# Intrinsic and extrinsic mechanisms alter neural cell fate specification in EPM1 Epilepsy

**DOI:** 10.1101/2024.11.20.624470

**Authors:** Andrea Forero, Veronica Pravata, Fabrizia Pipicelli, Alessandro Soloperto, Francesco Di Matteo, Zagorka Bekjarova, Elisa Frenna, Laura Canafoglia, Francesca Ragona, Giuseppina Maccarrone, Mariano Gonzalez Pisfil, Christian Whal-Schott, Filippo M. Cernilogar, Matthias Eder, Rossella Di Giaimo, Silvia Cappello

## Abstract

The extracellular milieu, including extracellular vesicles (EVs), plays a pivotal role in brain development by regulating neural processes such as proliferation, differentiation, and migration. In this study, we sought to elucidate the pathogenesis of progressive myoclonus epilepsy type 1 (EPM1), a disease caused by mutations in the *CSTB* gene, using cerebral organoids (COs) derived from EPM1 patient cells. The results demonstrate that EPM1 COs display increased electrophysiological activity and a disrupted excitatory/inhibitory balance. Single-cell RNA sequencing (scRNA-seq) analysis of ventral EPM1-Cos revealed an abnormal specification of progenitor fate, with a shift toward dorsal neuron identities at the expense of inhibitory interneurons. In addition, pathological alterations in EV biogenesis and cargo were identified, including aberrant Sonic Hedgehog (SHH) signaling, which may disrupt cortical patterning. These findings suggest that both intrinsic progenitor identity shifts and extrinsic EV-mediated signaling contribute to EPM1 pathology. Our study highlights potential therapeutic strategies mediated by EVs as a novel approach to mitigate disease progression.

## Introduction

The extracellular space, encompassing the extracellular matrix (ECM), soluble molecules and extracellular vesicles (EVs), is critical for brain development, influencing cellular processes such as proliferation, differentiation and migration (Forero et al., 2024; Long & Huttner, 2022). A clear example of its importance is observed during cortical development, where both excitatory projection neurons (ENs) and inhibitory interneurons (INs) rely on cues from the extracellular environment to reach their proper locations within the cerebral cortex. ENs migrate radially from the dorsal pallium into the cortical plate, while INs are generated in the ventral telencephalon and migrate tangentially to the neocortex (Marín & Rubenstein, 2001; Silva et al., 2019; Wichterle et al., 1999). Their migration is directed by a complex network of extracellular signals, including molecules and proteins secreted into the extracellular space via EVs (Amin & Borrell, 2020; Forero et al., 2024; Pipicelli et al., 2023). Disruptions within the extracellular space can lead to significant neuronal dysfunction (Long & Huttner, 2019), which may interfere with proper neurodevelopment and contribute to a range of neurological conditions, including autism spectrum disorders and intellectual disabilities (Klingler et al., 2021). Progressive Myoclonus Epilepsy Type I (EPM1), while not directly classified as a neurodevelopmental disorder, shares common features of neuronal dysfunction often observed in such disorders. It is the most prevalent type of progressive myoclonus epilepsy (PME), a group of inherited disorders marked by myoclonus, epilepsy, and progressive neurological decline (Berkovic et al., 1986). EPM1 is inherited in an autosomal recessive manner and is primarily caused by biallelic mutations in the *CSTB* gene, which encodes Cystatin-B, a ubiquitous 98-amino-acid protein (Lalioti et al., 1997).The majority of EPM1 patients (∼90%) carry a 12-nucleotide repeat expansion mutation in the promoter region of the *CSTB* gene, leading to a drastic reduction in CSTB transcript and protein levels, with levels reduced to less than 10% of normal (Joensuu et al., 2007). The clinical manifestations of EPM1 vary based on the genetic mutations. Patients with compound heterozygous mutations, involving the repeat expansion and a truncating null mutation, exhibit a severe form of the disease, characterized by earlier onset and accelerated progression (Canafoglia et al., 2012; Koskenkorva et al., 2011). Homozygous patients with two null mutations present the most severe form, neonatal-onset progressive encephalopathy (Mancini et al., 2016; O’Brien et al., 2017). These variations highlight the crucial role of CSTB in normal brain development. CSTB was initially identified as an inhibitor of cysteine proteases, essential for the regulation of cellular proteostasis by protecting against proteases leaking from lysosomes (Turk & Bode, 1991). Beyond this, CSTB protein has a proposed chaperone-like function (Rispoli et al., 2013; Žerovnik, 2022), and is also involved in various cellular processes, including modulating chromatin-associated proteolytic activity during neurogenesis (Daura et al., 2022), interacting with cytoplasmic proteins related to growth and differentiation (Di Giaimo et al., 2002), mTOR signaling (Trstenjak-Prebanda et al., 2023), and autophagy (Polajnar et al., 2014; Žerovnik, 2024). CSTB also plays a significant role in synaptic plasticity (Di Giaimo et al., 2020; Gorski et al., 2023; Penna et al., 2019; Pizzella et al., 2024). In CSTB knockout (*Cstb*^*−/−*^) mice, which serve as a model for EPM1, early gene expression alterations in the cerebellum and granule neurons have been associated with early synaptic changes and inflammation (Joensuu et al., 2014). These alterations suggest disrupted synaptic maturation, with a particular emphasis on GABAergic signaling. Impaired GABAergic inhibition has been documented in EPM1 patients (Buzzi et al., 2012; Silvennoinen et al., 2023), who also exhibit EEG abnormalities and progressive cerebral and cerebellar atrophy (Canafoglia et al., 2004; Joensuu et al., 2014; Kyllerman et al., 1991; Singh & Hämäläinen, 2024). This underscores CSTB’s crucial role in maintaining neural function. Our previous research demonstrated that CSTB is not only essential for intracellular regulation but also secreted to instruct neighboring cells during neurogenesis (Di Matteo et al., 2020). Functional CSTB levels influence the expansion of progenitor cells both intrinsically and extrinsically, and pathologically low CSTB leads to reduced recruitment of interneurons while driving premature differentiation of progenitors (Di Matteo et al., 2020). Together, these findings highlight the multifaceted roles of CSTB in neural development and function.

Given that IN migration is altered in EPM1 and considering CSTB’s role in extracellular signaling, we hypothesize that these alterations could extend beyond migration, potentially affecting interneuron development itself. Aberrant CSTB signaling may therefore lead to broader defects in IN maturation and function, contributing to the overall pathology of EPM1. In this study, we employ human cerebral organoids (COs) generated from EPM1 patient lines (EPM1-COs) to model the disease. We show that EPM1-COs can recapitulate the pathologically altered electrophysiological properties observed in EPM1 patients. We then focus on ventrally patterned COs, which give rise to INs, and demonstrate that EPM1 ventral organoids exhibit altered patterning, leading to an impaired excitatory/inhibitory balance. Importantly, we propose that CSTB alters the extracellular environment through changes in the biogenesis and composition of EVs, particularly by modulating SHH levels within these vesicles. This altered SHH signaling via EVs is likely a critical factor in influencing neural differentiation and fate, thereby playing a significant role in the pathogenesis of EPM1.

## Results

### Physiological alteration of neuronal activity in EPM1-COs is caused by changes in the neuronal fate

To investigate the mechanisms underlying the increased neuronal excitability and disrupted excitatory-inhibitory balance observed in EPM1, we generated cerebral organoids (COs) from two patient and two control induced Pluripotent Stem Cell (iPSC) lines and cultured them for 8 months to probe their electrophysiological properties and assess whether this model could recapitulate the patient phenotype. We performed silicon probe extracellular recordings to reliably record from spontaneously firing neurons at several locations within a CO as previously shown (Di Matteo et al, 2024). Our findings revealed that 8-month-old EPM1-COs exhibit significantly higher electrophysiological activity compared to two control lines (CTRL-COs), as indicated by spike distribution and average spike number (Fig. 1A-D; Appendix Fig. S1A-F). Specifically, EPM1-COs showed a greater percentage of high-frequency spikes, defined as spikes with an inter-spike interval (ISI) below 500ms, compared to CTRL-COs (Fig. 1D; Appendix Fig. S1C). The observed hyperactivity in EPM1-COs may result from an imbalance between excitatory and inhibitory neurons, which is a common feature in epileptic conditions (Sohal & Rubenstein, 2019). To investigate this possibility, we examined the differentiation fate of unpatterned neural progenitor cells (NPCs) derived from patient or control cells. After eight weeks of differentiation, we found a significant increase in the number of excitatory marker-expressing neurons in EPM1 cultures, along with a corresponding decrease in inhibitory marker-expressing neurons compared to controls (Fig. 1E-H and Appendix Fig. S1E-F). These findings suggest that the hyperactivity observed in EPM1-COs could be driven by an imbalance in the production of neuronal subtypes, with a shift favoring excitatory over inhibitory neurons. To further validate these results, we generated neurons derived from both ventral (vNeurons) and dorsal (dNeurons) progenitors, dissociated from 30-day-old patterned EPM1- and CTRL-COs. Dorsally regionalized COs (dCOs) mimic the cellular architecture of the dorsal telencephalon, resulting in an enrichment of excitatory neurons. In contrast, ventrally regionalized COs (vCOs) resemble the ganglionic eminences, which predominantly give rise to inhibitory interneuron populations (Bagley et al., 2017). This approach allows us to directly compare the distinct neuronal subtypes influenced by regional patterning, providing insights into how EPM1 impacts both excitatory and inhibitory specification. In line with the previous results, vNeurons from ventrally patterned EPM1 cerebral organoids (EPM1-vCOs) exhibited a different proportion of GAD67-positive (inhibitory) and TBR1-positive (excitatory) neurons compared to controls (Fig. 1I, J, L, M; Appendix Fig. S1G). Specifically, ventral EPM1 progenitors yielded fewer GAD67-positive neurons and more TBR1-positive neurons than controls, while dorsally derived progenitors did not exhibit significant differences in neuronal specification when compared to control NPCs (Fig. 1I, K, L, N; Appendix Fig. S1H).

**Fig. 1:**
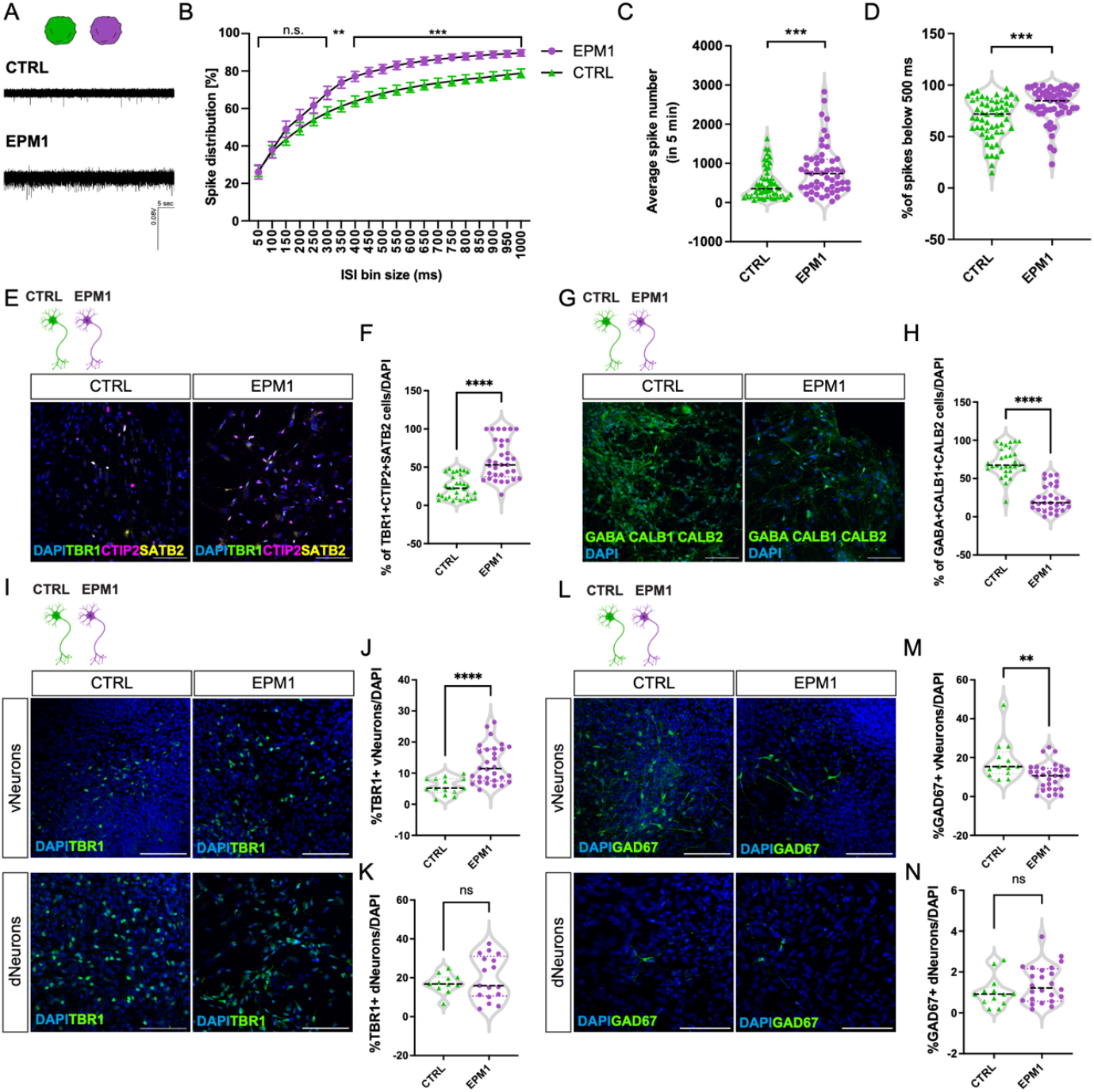
Altered electrophysiological activity and neuronal composition in EPM1. (A) Representative electrophysiological traces of CTRL and EPM1 cerebral organoids (COs). (B) Percentage of spike distribution at interspike interval bins ranging from 50 to 1000 ms for CTRL and EPM1 COs. n = 53-55 per condition. Statistical significance was determined using a Mann-Whitney test, **p<0.01, ***p<0.001. (C) Average spike number detected within 5 min for CTRL and EPM1 COs. n = 53-55 per condition. Statistical significance was determined using the a Mann-Whitney test, ***p<0.001. (D) Percentage of spikes below 500 ms detected in CTRL and EPM1 COs. n = 53-55 per condition. Statistical significance was determined using the Mann-Whitney test, ***p<0.001. (E) Representative images of CTRL and EPM1 neurons immunostained for excitatory neuronal markers TBR1 (Green), CTIP2 (Magenta) and SATB2 (Yellow). Nuclei were makers with DAPI (Blue). Scale bar: 100 µm. (F) Quantification of the percentage of cells positive for TBR1/CTIP2/SATB2 in CTRL versus EPM1 neural cultures normalized by total DAPI nuclear staining. n = 30 per condition. Statistical significance was determined using the Mann-Whitney test, ****p<0.0001. (G) Representative images of CTRL and EPM1 neurons immunostained for inhibitory neuronal markers GABA/Calretinin (CALB2)/Calbindin (CALB1). Nuclei were makers with DAPI (Blue). Scale bar: 100 µm. (H) Quantification of the percentage of cells positive for GABA/ CALB2/ CALB1 in CTRL versus EPM1 neural cultures. n = 33 per condition. Statistical significance was determined using the unpaired Student’s t-test, ****p<0.0001. (I) Representative images of TBR1-positive cells in CTRL and EPM1 neurons derived from dorsal (dNeurons) and ventral (vNeurons) progenitors. Nuclei were makers with DAPI (Blue). Scale bar: 100 µm. (J) Quantification of TBR1-positive cells/DAPI in CTRL versus EPM1 vNeurons. n = 15-30 per condition. Statistical significance was determined using the Mann-Whitney test, ****p<0.0001. (K) Quantification of TBR1-positive cells/DAPI in CTRL versus EPM1 dNeurons. n = 11-17 per condition. Statistical significance was determined using the unpaired Student’s t-test, n.s. not significant. (L) Representative images of GAD67-positive cells in CTRL and EPM1 neurons derived from dorsal (dNeurons) and ventral (vNeurons) progenitors. Nuclei were makers with DAPI (Blue). Scale bar: 100 µm. (M) Quantification of GAD67-positive cells/DAPI in CTRL versus EPM1 vNeurons. n = 13-31 per condition. Statistical significance was determined using the unpaired Student’s t-test, **p<0.01. (N) Quantification of GAD67-positive cells/DAPI in CTRL versus EPM1 dNeurons. n = 13-24 per condition. Statistical significance was determined using the unpaired Student’s t-test, n.s. not significant. Abbreviations: milliseconds (ms); minutes (min).

To assess whether EPM1 vNeurons exhibited heightened neuronal activity driven by the altered differentiation of ventral progenitors, we employed Multi Electrode Arrays (MEA) analysis. For MEA recordings, we focused on the channel with the highest activity to record the spike activity and investigate potential alterations in the neuronal network properties. The analysis confirmed heightened neuronal activity in EPM1 patient-derived cultures after 10 weeks of differentiation (Appendix Fig. S1I-J). When combining dorsal control cells with ventral EPM1 cells, EPM1 vNeurons exhibited markedly higher activity than their control counterparts (Appendix Fig. S1K-L). These findings suggest that ventral NPCs from EPM1 patients undergo an aberrant differentiation pathway, contributing to the altered electrophysiological properties observed in EPM1-COs.

Overall, these results suggest that the altered excitatory-inhibitory balance and increased neuronal activity in EPM1 -COs may be associated with the aberrant differentiation of ventral progenitors, leading to an imbalance that drives increased excitability in the neuronal network.

### EPM1 ventral progenitors and neurons change cell identity

Building on our observations of altered neuronal activity and disrupted cell fate decisions in EPM1-derived cells, we sought to further investigate the molecular mechanisms underlying these changes by analyzing the transcriptomic profiles of progenitors and their neuronal progeny. To this end, we performed single-cell RNA sequencing (scRNA-seq) on 40-day-old vCOs derived from EPM1 (EPM1-vCOs) and control (vCOs) lines. We selected day 40 as a critical time point for exploring the effects of CSTB protein dysregulation on neuronal differentiation and maturation. *CSTB* mRNA becomes detectable as early as day 16, but CSTB protein levels only emerge by day 40 (Di Matteo et al., 2020). Following stringent quality control, we retained 10,972 high-quality cells across EPM1 and control ventral organoids. To visualize cellular heterogeneity, we integrated cells from vCOs (two control lines) and EPM1-vCOs (two patient lines), projecting them onto a two-dimensional UMAP space (Fig. 2A, Appendix Fig. S2A). The resulting clusters encompassed a wide array of cell types, including neural crest cells, neural progenitors (e.g., forebrain and striatal radial glia), and mature neurons. This dataset enabled us to interrogate the transcriptional similarities and differences between progenitors and neurons from both control and EPM1 origins.

**Fig. 2.**
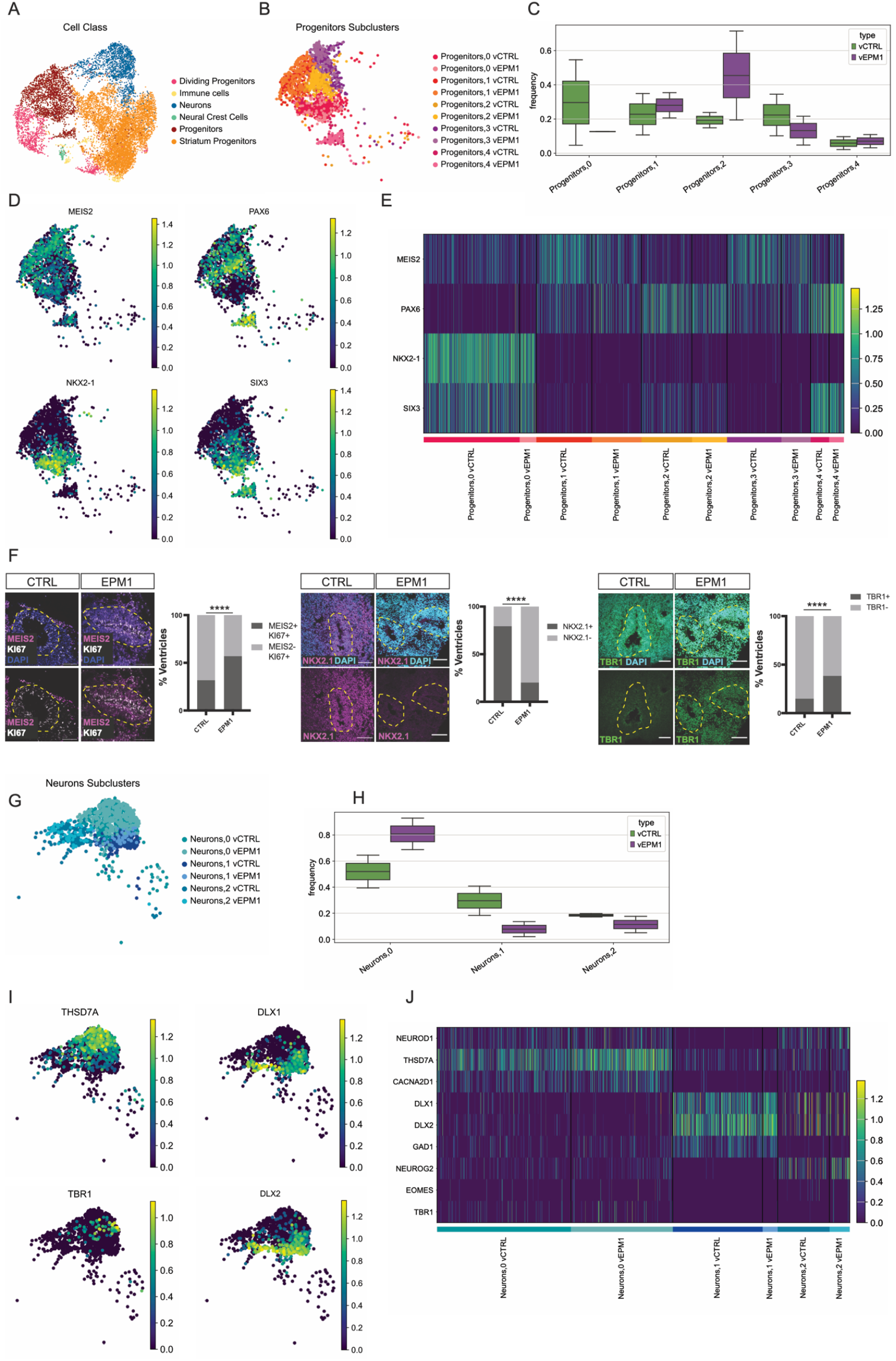
Changes in Progenitor and Neuron Identity in Ventral EPM1 Cerebral Organoids. (A) UMAP visualization of scRNA-seq data from CTRL and EPM1 vCOs, clustered by cell types (Dividing progenitors, Immune cells, Neurons, Neural Crest cells, Progenitors, Striatum progenitors). (B) UMAP visualization of five subclusters within the Progenitors group (Progenitors 0, 1, 2, 3, and 4). Lighter colors represent Progenitors derived from vEPM1, while darker colors indicate those from vCTRL. (C) Box plot illustrating the frequency of vCTRL and vEPM1 cells in the five subclusters (Progenitors 0, 1, 2, 3, and 4) identified within the Progenitors cluster. (D) UMAP visualization showing spatial expression of MEIS2, PAX6, NKX2-1, and SIX3 across cells in the progenitor clusters. (E) Heatmap showing the expression levels of MEIS2, PAX6, NKX2-1, and SIX3 in the cells composing the five Progenitors subclusters (Progenitors 0, 1, 2, 3, and 4). (F) Representative images of CTRL and EPM1 vCOs immunostained for MEIS2, NKX2-1, KI67, and TBR1, and quantification of the percentage of ventricles positive for MEIS2/KI67, NKX2-1, and TBR1 in CTRL versus EPM1 vCOs. n = 9-11. Statistical significance was determined using an exact binomial test, ****p<0.0001. (G) UMAP visualization of the three Neurons subclusters (Neurons 0, 1, and 2) identified within the Neurons cluster. Lighter colors represent Neurons derived from vEPM1, while darker colors indicate those from vCTRL. (H) Box plot illustrating the frequency of vCTRL and vEPM1 cells in the Neurons subclusters (Neurons 0, 1, and 2). (I) UMAP visualization highlighting the expression of excitatory neuronal markers (THSD7A), inhibitory neuronal markers (DLX1, DLX2), and dorsal telencephalon markers (TBR1). (J) Heatmap showing the expression levels of NEUROD1, THSD7A, CACNA2D1, DLX1, DLX2, GAD1, NEUROG2, EOMES, and TBR1 in the cells composing the Neurons subclusters (Neurons 0, 1, and 2).

We focused initially on the diversity within the progenitor population. Progenitors expressed classical markers, including *SOX2, NES*, and *VIM* (Appendix Fig. S2B), and were further subdivided into five distinct subclusters: ‘Progenitor,0’ through ‘Progenitor,4’ (Fig. 2B). Notably, the frequencies of the Progenitor,0 and Progenitor,2 subclusters differed significantly between EPM1 and control cells (Fig. 2C, Appendix Fig. S2C), with Progenitor,0 being predominantly enriched in control cells, while Progenitor,2 was overrepresented in EPM1 cells. Interestingly, Progenitor,0 was characterized by the expression of medial ganglionic eminence (MGE) markers such as *NKX2-1* and *SIX3*, whereas Progenitor,2 exhibited dorsal progenitor marker *PAX6* expression (Fig. 2D-E). The Progenitor,1 and Progenitor,3 subclusters, meanwhile, displayed a combination of *MEIS2* and *PAX6*, markers commonly associated with the dorsal lateral ganglionic eminence (dLGE) (Fig. 2D-E). To corroborate our scRNA-seq findings, we performed immunofluorescence analysis, which confirmed a reduction in NKX2-1+ progenitors and an increase in MEIS2+ progenitors in EPM1-vCOs compared to control vCOs (Fig. 2F). Further validation through immunostaining of dissociated cells from EPM1-vCOs reinforced the predominance of MEIS2+ progenitors in the EPM1 condition (Appendix Fig. S2D).

Next, we characterized the neuronal populations, which expressed common neuronal markers such as *DCX* and *MAP2* (Appendix Fig. S2E). We identified three subclusters: ‘Neurons,0’, ‘Neurons,1’, and ‘Neurons,2’ (Fig. 2G). EPM1-derived cells were enriched in the Neurons,0 subcluster, while control-derived cells were predominantly present in the Neurons,1 subcluster (Fig. 2H, Appendix Fig. S2F). Consistent with the progenitor data, the Neurons,0 subcluster was enriched for excitatory neuron markers, including *THSD7A, CACNA2D1*, and *NEUROD1* (Fig. 2I-L, Appendix Fig. S2G), while Neurons,1, primarily populated by control cells, expressed genes associated with inhibitory neurons, such as *DLX1, DLX2*, and *GAD1* (Fig. 2I-L, Appendix Fig. S2G). Immunofluorescence on 70-day-old vCOs further confirmed an increase in TBR1+ excitatory neurons in EPM1-vCOs (Fig. 2F), whose expression is regulated by *NEUROG2* and the neurogenic transcription factor *EOMES* (Appendix Fig. S2G). Additionally, consistent with the increased expression of dLGE progenitor markers—known to give rise to medium spiny neurons—we observed an enrichment of DARPP32+ striatal neurons in EPM1-vCOs by immunostaining (Appendix Fig. S2H). These findings indicate that EPM1-vCOs exhibit altered patterning, with progenitors and neurons shifting from MGE to both dorsal and LGE fates, potentially contributing to the imbalance in excitatory and inhibitory neural populations.

### EPM1 cells exert a dominant non-cell-autonomous effect on control cells

Based on our previous findings that CSTB is secreted and can remodel the extracellular environment, thereby influencing surrounding cells (Di Matteo et al., 2020), we investigated whether altered levels of CSTB can affect cell fate in a non-cell-autonomous manner. We generated mosaic vCOs (mvCOs) containing GFP-labeled CTRL cells (mvCTRL) and non-labeled EPM1 cells (mvEPM1). Single-cell RNA sequencing was performed on FACS sorted cells from 40-day-old mvCOs. The datasets from mvCTRL and mvEPM1 cells were integrated and cell diversity mapped with UMAP (Fig. 3A). The identified cell types spanned dividing progenitors, progenitors, neurons and immune cells.

**Fig. 3.**
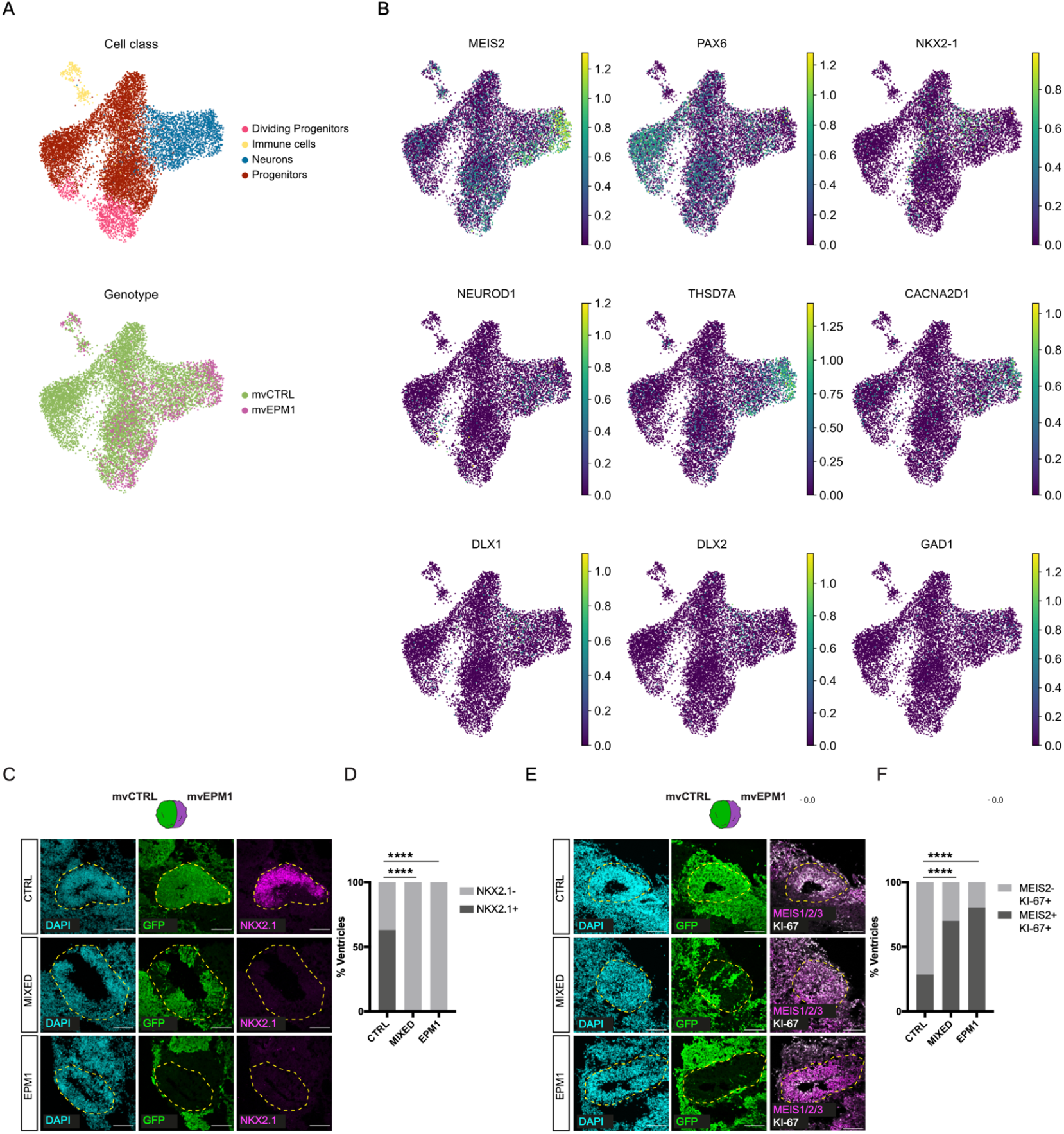
Cell-non-autonomous effect of EPM1 cells in mosaic vCOs. (A) UMAP visualization of scRNA-seq data of mosaic CTRL (mvCOs) and EPM1 (mvEPM1) vCOs, clustered by cell classes (top; Dividing progenitors, Immune cells, Neurons, Progenitors) and genotype (bottom; mvCTRL and mvEPM1). (B) UMAP visualization of the expression of progenitor markers (MEIS2, PAX6, NKX2-1), excitatory neuronal (NEUROD1, THSD7A, CACNA2D1), and inhibitory neuronal (DLX1, DLX2, GAD1) markers. (C) Representative images of CTRL (GFP+), EPM1 (GFP-), and mixed (GFP+/GFP-) ventricles in mvCOs, immunostained for NKX2-1. (D) Quantification of percentage of CTRL (GFP+), EPM1 (GFP-), and mixed (GFP+/GFP-) ventricles positive for NKX2-1. n = 44. Statistical significance was determined using an exact binomial test, ****p<0.0001. (E) Representative images of CTRL (GFP+), EPM1 (GFP-), and mixed (GFP+/GFP-) ventricles in mvCOs, immunostained for MEIS1/2/3 and KI67. Quantification of percentage of CTRL (GFP+), EPM1 (GFP-), and mixed (GFP+/GFP-) ventricles positive for MEIS1/2/3 and KI67. n = 26. Statistical significance was determined using an exact binomial test, ****p<0.0001.

Strikingly scRNAseq data indicated that mvCTRL cells in close contact with mvEPM1 cells acquired features of the vEPM1 cells during neurodevelopment. Notably, the expression of the MGE progenitor marker NKX2.1 was dramatically decreased in cells from mvCOs compared to cells from vCOs regardless of their genotype, while the expression of LGE markers such as MEIS2 increased (Fig. 3B and Fig. 2D). Accordingly, the neuronal output of mvCTRL cells was strongly affected by the pathological environment, with an increased expression of excitatory markers and a reduced expression of inhibitory markers (Fig 3B and Appendix S3A). mvCTRL (GFP+), mvEPM1 (GFP-) and mixed (GFP+/-) ventricles were then analyzed by immunofluorescence on mvCOs at day 30. While CTRL ventricles contained NKX2.1+cells, the germinal zone of the mixed ventricles showed an identity closer to EPM1 ventricles, with a decreased number of NKX2.1+ ventricles and an increased number of MEIS2+ ventricles (Fig. 3C-F). Our results indicate that the onset of EPM1 relies on extrinsic mechanisms that can influence neurogenesis during development.

### EPM1 EVs present altered biogenesis, dynamics and composition

CSTB has been detected in the culture medium of COs, as previously reported (Di Matteo et al., 2020), and notably, has also been observed to be released via extracellular vesicles (EVs) (Forero et al., 2024; Pizzella et al., 2024). To explore the extrinsic mechanisms underlying EPM1, we investigated how EVs could alter the extracellular environment and influence neighboring cells. We isolated a mixed population of small EVs, including both exosomes and small microvesicles (100-300 nm, Appendix Fig. S4A), from the culture medium of vCOs and EPM1-vCOs using differential ultracentrifugation (Forero et al., 2024; Mathieu et al., 2019; Théry et al., 2018). Immuno-electron microscopy confirmed the presence of CSTB within EVs, with some EVs also positive for the EV marker CD81 (Fig. 4A). Interestingly, CSTB abundance in EVs changes during CO development with 2 peaks at d40 and at later stages of development together with other developmental markers like RELN and MAP2 (FIG. 4B, Forero et al., 2024). Moreover, CSTB overlaps with CST3 in EVs at later stages of development (Appendix Fig. S4B) and, interestingly, CST3 has a beneficial effect in mice lacking CSTB (Kaur et al., 2010).

**Fig. 4.**
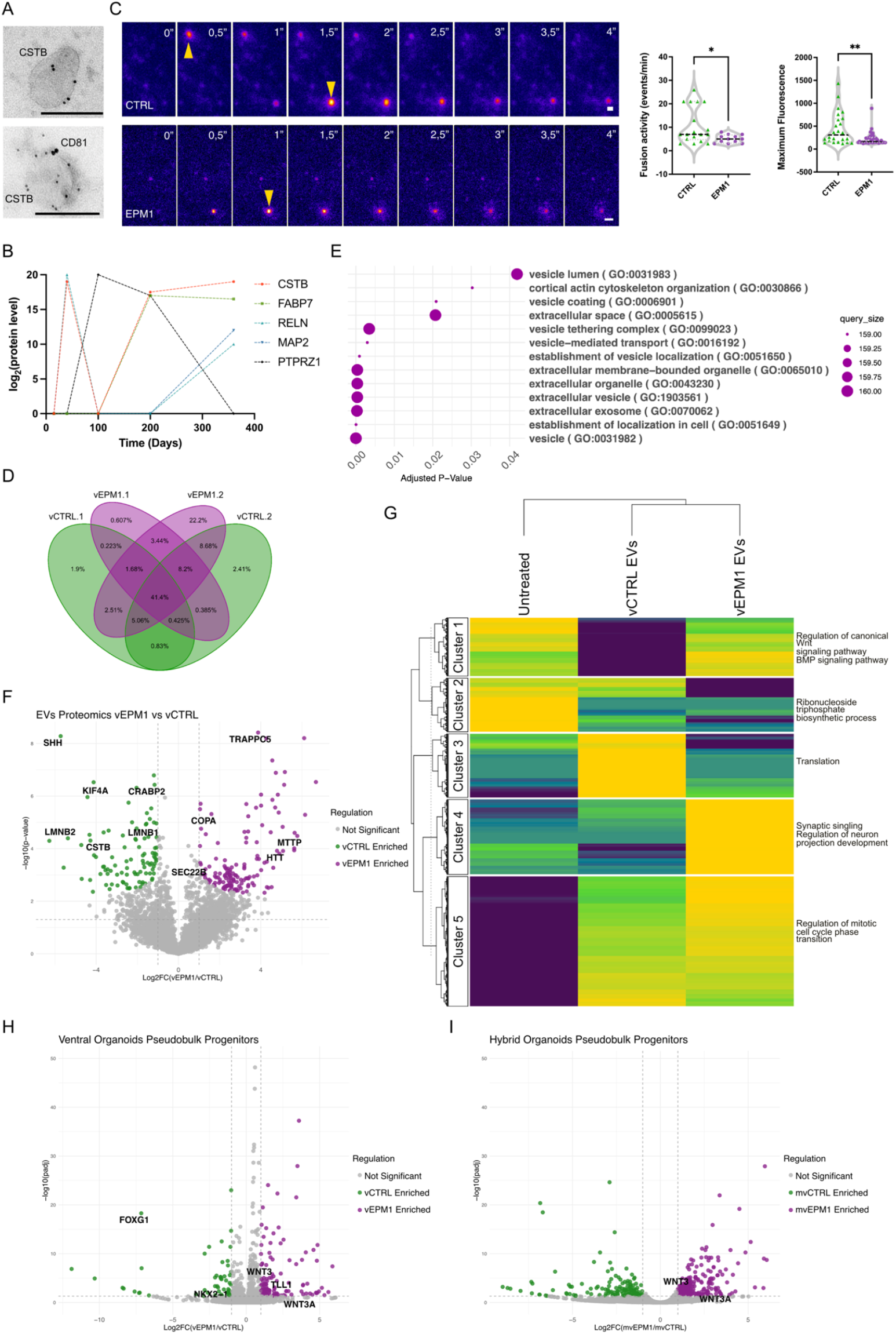
Alterations in EV biogenesis and composition in EPM1. (A) Immuno-electron micrographs of CD81 (large dots) and CSTB (small dots) in EVs collected from COs. Scale bar: 1 µm. (B) Expression levels of CSTB, FABP7, RELN, MAP2 and PTPRZ1 protein levels in extracellular vesicles derived from COs at different developmental stages (expressed in days). (C) Time lapse images of MVB-plasma membrane fusion events (arrow heads) in CTRL and EPM1 neurons (left). Quantification of the fusion activity (events/min) and Maximum Fluorescence of the fusion events detected in CTRL and EPM1 neurons. Every dot refers to a single cell analyzed, n= 29 (CTRL), n= 26 (EPM1). Statistical significance was determined using the Mann-Whitney U test, *p<0.05, **p<0.01. Scale bar: 1 µm. (D) Venn diagram depicting the percentage of unique and common proteins detected in EVs from vCOs from two control iPSC lines (vCTRL.1 and vCTRL.2) and two EPM1 patient iPSCS lines (vEPM1.1, vEPM1.2). (E) Gene Ontology (GO) enrichment analysis of the unique proteins detected in EPM1-vEVs. (F) Volcano plot of proteins enriched in CTRL-vEVs (vCTRL, green) versus EPM1-vEVs (vEPM1, magenta). (G) Clustered heatmap showing the expression levels of the differentially regulated genes in NPCs following the acute treatment with vEVs and vEPM1-EVs or no treatment. GO terms enriched in each cluster are listed on the right. (H) Volcano plot showing pseudobulk analysis for the Progenitor cluster of scRNA-seq data from ventral hybrid COs. Genes enriched in CTRL are highlighted in green, and genes enriched in EPM1 are highlighted in magenta. (I) Volcano plot showing pseudobulk analysis for the Progenitor cluster of scRNA-seq data from ventral COs. Genes enriched in CTRL are highlighted in green, and genes enriched in EPM1 are highlighted in magenta.

Given that CSTB is secreted in EVs and we identified non-cell-autonomous effects on excitatory/inhibitory differentiation in mosaic vCOs, we hypothesized that EVs could be key players in this phenotype. To explore this, we next aimed to investigate the biogenesis and secretion dynamics of EVs. We began by employing Nanoparticle Tracking Analysis (NTA), a real-time technique used to measure the size and concentration of nanoparticles, such as EVs, by tracking their Brownian motion. NTA was employed to monitor EV secretion during the early stages of differentiation of NPCs into neurons. EPM1 young neurons (d5 to d12 of differentiation) secreted a lower amount of EVs with an average diameter of 115-165nm compared to CTRL cells and showed a trend toward larger diameter vesicles (Appendix Fig. S4C). These findings prompted us to further examine EV secretion dynamics under EPM1 conditions. To achieve this, we employed live imaging of membrane fusion activity in both control and EPM1 neurons using TIRF microscopy. This was performed after overexpressing CD63-pHluorin, a tetraspanin-based pH-sensitive optical reporter that detects multivesicular body–plasma membrane (MVB-PM) fusion events (Verweij et al., 2018) (Fig. 4C). EPM1 neurons displayed a decreased number of fusion events per minute, indicating altered EV secretion in pathological conditions.

After identifying differences in EV secretion, we next investigated whether the protein cargo of EVs differed between control and EPM1 conditions. Using mass spectrometry, we profiled the protein content of EVs from CTRL (vEVs) and EPM1 (EPM1-vEVs), detecting 2,690 proteins for vEV1, 3,362 for vEV2, 2,805 for EPM1-vEV1, and 4,654 for EPM1-vEV2. Consistent with previous studies (Théry et al., 2018), standard EV markers (CD9, CD63, CD81, CD82, PDCD6IP, and TSG101) were present in all samples (Appendix Fig. S4D), while negative markers such as ALB and AGO (Théry et al., 2018) were not detectable. We compared proteins secreted by each control and patient line to identify EPM1-specific vEV signatures, finding that control and EPM1-vEVs shared a large proportion of proteins (41.4%) (Fig 4D). However, 3.4% of the total detected proteins (170) were unique to EPM1-vEVs (Fig. 4D) and enriched for processes involved in cell trafficking from the ER to the extracellular space (Fig.4E). Among the proteins unique to vEVs (0.8%, 41 proteins), no specific Gene Ontology (GO) terms were identified.

We then focused on differentially expressed proteins (DEP) between vEVs and EPM1-vEVs (Fig.4F). GO term analysis of DEPs confirmed the accumulation of proteins involved in cell trafficking within EPM1-vEVs (Appendix Fig. S4E). In contrast, proteins depleted in EPM1-vEVs were associated with altered processes related to mRNA processing, nuclear pore localization, mitotic spindle formation, cytoskeletal organization, and axogenesis, all of which suggest pathological changes in gene expression, cell division, and neuronal maturation (Appendix Fig. S4E). As expected, CSTB was significantly downregulated in EPM1-vEVs.

Notably, many proteins altered in EPM1-vEVs are encoded by genes associated with human brain diseases, according to Disgenet (www.disgenet.com, Appendix Fig. S4F-G). Among the DEPs in EPM1-vEVs, we identified associations with epilepsy, intellectual disabilities, and brain malformations, underscoring the critical role of EV-mediated signaling in neurodevelopmental disorders.

Among the differentially expressed proteins, SHH was significantly enriched in vEVs and emerged as a promising candidate (Fig. 4F). Notably, we previously identified SHH in EVs derived from ventrally regionalized organoids (Forero et al., 2024), further supporting its role in these structures. In the forebrain, SHH is primarily secreted from the prechordal plate and the ventral midline, including the notochord and floor plate. This signaling establishes a gradient that specifies the identity of neural progenitor cells along the dorsoventral axis (Ericson et al., 1995; Puelles & Rubenstein, 2003; Rallu et al., 2002; Q. Xu et al., 2005).

Based on these findings, we hypothesized that the observed differences in EV composition could potentially explain the altered differentiation outcomes in progenitor identity. To test this hypothesis, we explored both short- and long-term effects of EVs on progenitor cells. For short-term effects, we treated NPCs with CTRL vEVs or EPM1-vEVs for 12 hours and analyzed their transcriptome through bulk RNA sequencing (Fig.4G). Compared to untreated NPCs, vEVs induced several transcriptomic changes, confirming the role of EVs in rapid cellular responses. Moreover, transcriptomic differences between vEV and EPM1-vEV-treated NPCs suggested that the altered protein cargo of EPM1-vEVs can indeed influence NPC transcripts quickly. Although the short-term exposure to EVs did not result in significant effects on the SHH pathway, we identified several changes associated with cell trafficking, as well as the regulation of WNT and BMP signaling pathways (Fig.4G). This aligns with the understanding that lower levels of SHH allow the development of more dorsal structures by limiting the expression of ventralizing factors and permitting the action of dorsalizing signals like BMPs and WNTs. These findings suggest that EPM1-vEVs can impact the overall trafficking machinery, EV biogenesis, and contribute to the regulation of key signaling pathways involved in dorsal-ventral patterning.

We then examined long-term effects by comparing DEG between EPM1 and CTRL cells derived from vCOs or mvCOs, using single-cell RNA sequencing data. We performed pseudobulk RNA analyses to investigate whether general transcriptional differences could be attributed to non-cell-autonomous effect. In the vCOs we identified *NKX2*.*1* enriched in vCTRL progenitors while *WNT3* and *WNT3A* were enriched in vEPM1 progenitors (Fig. 4H). On the contrary, when the pseudobulk analysis was performed on progenitors derived from mosaic organoids, *NKX2*.*1* was no longer amongst the DEG (FIG4I). As high concentrations of SHH promote the activation of transcription factors like *NKX2*.*1* and *OLIG2* (Q. Xu et al., 2005), this indicates that pathological non-cell-autonomous levels of SHH could alter MGE progenitor fate.

## Discussion

In this study, we aimed to explore the mechanisms through which both intrinsic and extrinsic factors contribute to EPM1 disease progression. Using brain organoids derived from iPSCs from EPM1 patients harboring mutations in the *CSTB* gene, we successfully recapitulated impaired electrophysiological activity, highlighting defects in the excitatory/inhibitory balance that likely contributes to the onset of epilepsy and other neurological symptoms. Previous studies have documented reduced GABAergic inhibition and cortical thinning in EPM1 patients and *CSTB* knockout mouse models (Buzzi et al., 2012; Silvennoinen et al., 2023). Here we showed that low levels of CSTB in EPM1 result in intrinsic effects such as progenitor cell misspecification, and extrinsic effects, mediated by non-cell autonomous cues conveyed by EVs, which influence the neighboring cells. We propose that reduced GABAergic signaling depends on the pathological misspecification of progenitor cell fate, which led to an altered neuronal output characterized by an unexpected shift toward dorsal cortex and dLGE identities. Cell type-specific alterations in the ventral fate of progenitor cells during early development leading to fewer GABAergic neurons have also been described in other neurodegenerative disorders like Huntington’s disease (Galimberti et al., 2024). These common pathological features underscore the importance of investigating aspects of neurodevelopment in diseases not directly classified as a neurodevelopmental disorders.

Importantly, our study expands upon these findings by showing that these alterations arise from both intrinsic factors, and extrinsic influences, particularly through EVs, which can modulate neurogenic niche.

Interestingly, the unique protein content of EPM1-vEVs suggests a link between altered trafficking mechanisms and disease pathology. In EPM1 EVs, we identified many components of the TRAPP complex, which regulates vesicle tethering and fusion (Zhao et al., 2017). Disruptions in TRAPP function may impair EV production and release, affecting cell communication in the developing brain, leading to defects in the delivery of proteins and lipids necessary for membrane biogenesis and vesicle formation. To further support our data, we demonstrated that EPM1 cells release fewer CD63+ EVs, which supports the disruption in the biogenesis and release of EVs. Overall, these data support the idea that EPM1 cells may have defects in the late stages of EV biogenesis or in the efficient release of these vesicles.

Among the proteins altered in EPM1 EVs, we identified sonic hedgehog (SHH), a key player in dorsoventral patterning. This highlights a potential mechanism by which EV-mediated signaling may contribute to the altered cortical patterning observed in EPM1. Low levels of SHH in EPM1-vEVs could drive progenitor cell fate changes. Indeed, human COs generated with a SHH protein gradient clearly show that dorsal (PAX6+, TBR2+, TBR1+) cerebral cortex-like tissue and DARPP32+ striatum-like tissue emerge in areas distal from the SHH source tissue, while MGE-like tissue is detected in proximal areas (Cederquist et al., 2019). This finding highlights the importance of progenitor identity in maintaining cortical E/I balance and suggests that disruptions in this process may be a key driver of EPM1 pathology.

Although control vEVs contain SHH, the absence of this signaling in EPM1 cells may disrupt the gradient necessary for proper patterning, leading to a dominance of the pathological signals from EPM1-vEVs. This suggests that the SHH gradient is essential for maintaining normal cortical development, and its disruption in EPM1 could contribute to the observed phenotypic alterations. The observed changes in EV content, suggest that EVs may play a crucial role in progenitor specification and the subsequent disruption of cortical patterning in EPM1. This is consistent with recent findings by Pipicelli et al. (2023), who reported that pathological EVs secreted by vCOs with mutations in the ECM component *LGALS3BP* gene influence progenitor cell fate and dorso/ventral patterning, further highlighting the role of EVs in cell fate specification during brain development.

In conclusion, our study reveals both intrinsic and extrinsic mechanisms contributing to the pathogenesis of EPM1, with significant implications for understanding the disease and developing potential therapeutic strategies. The altered progenitor fate and the role of pathological EVs in disrupting cortical development highlight the complexity of EPM1 and underscores the need for continued research into the cellular and molecular underpinnings of this and related disorders.

## Materials and Methods

### IPSCs culture

iPSCs were previously reprogrammed from 2 control lines of fibroblasts (Klaus et al., n.d.) and PBMCs origin (Di Matteo et al., 2020) and from 2 EPM1 patient lines (Di Matteo et al., 2020). iPSCs were cultured on Matrigel (Corning) coated plates (Thermo Fisher, Waltham, MA, USA) in mTesR1 basic medium supplemented with 1x mTesR1 supplement (Stem Cell Technologies, Vancouver, Canada) at 37°C, 5% CO2 and ambient oxygen level. Passaging was done using accutase (Stem Cell Technologies) treatment.

### Generation of labelled iPSC lines

The GFP-labeled iPSC lines were generated using the piggyBac transposase (1ug) and PB-GFP (1ug) nucleofection (Chen & LoTurco, 2012). Single cells of iPSCs were transfected with the Amaxa Nucleofector 2b (program B-016). RFP positive colonies were picked and cultured on Matrigel (Corning/VWR International, 354234) coated plates in mTeSR1 basic medium (Stem Cell Technologies, 85850) supplemented with 1× mTeSR1 supplement (Stem Cell Technologies, 85850) at 37°C and 5% CO2.

### Generation of patterned human organoids

Patterned human organoids were generated according to (Bagley et al., 2017). Embryoid bodies (EBs) generated from iPSCs were patterned to have ventral and dorsal identity. During the neuronal induction step, EBs were treated individually with SAG (1:10,000) (Millipore, 566660) + IWP-2 (1:2,000) (Sigma-Aldrich, I0536) for inducing ventral identity, s with cyclopamine A (1:500) (Calbiochem, 239803) for inducing dorsal identity. After this point, the generation of organoids followed methods according to (Lancaster et al., 2013; Lancaster & Knoblich, 2014). Organoids were kept in 10-cm dishes on a shaker at 37°C, 5% CO2 and ambient oxygen level with medium changes every 3–4 days.

### Generation of mosaic human organoids (vmCO)

iPSCs from GFP-labeled iPSC control line and from EPM1 iPSCs, were dissociated into single cells using Accutase (Sigma-Aldrich, A6964), mixed together in a ratio of 30% GFP-control cells : 70% EPM1 cells and transferred in a total of approximately 9000 cells to one well of an ultra-low-attachment 96-well plate (Corning). The protocols continued as described in “Generation of patterned human organoids”.

### NPC and neuron generation and culture

Neural progenitor cells (NPCs) were generated and maintained as previously described (Zink et al., 2021). NPCs were generated from two control iPSC lines, which generated a ratio of 60% neurons and 40% astrocytes in accordance with this protocol, providing electrophysiologically mature neurons in a more physiological environment. Neural differentiation was conducted as previously described (Gunhanlar et al., 2018). All the cells were kept in an incubator at 37°C, 5% CO2 and ambient oxygen level with medium changes every 2–3 days.

### EV collection and analysis

In accordance with Théry et al., (Théry et al., 2018) and Forero et al. 2024 (Forero et al., 2024), EVs were collected from conditioned media from COs and 2D cultured cells by the following steps: centrifugation at 300g for 10 mins, supernatant centrifugation at 2000g for 10 mins at 4 °C, supernatant centrifugation at 10.000g for 30 mins at 4 °C, supernatant centrifugation at 100.000g for 90 mins at 4 °C in a fixed-angle rotor (TH865, Thermo Fisher Scientific), followed by pellet wash with 1x PBS and centrifugation at 100.000g for 90 mins at 4 °C. Alternatively, miRCURY Exosome Cell/Urine/CSF Kit (Qiagen, 76743) was used to isolate EVs from conditioned medium according to the manufacturer instructions. For NPCs, EVs were collected from three independent cultures of control NPCs. Neuronal EVs were collected from three independent neuronal differentiation cultures. Similarly, astrocyte EVs were collected from three independent cultures of astrocytes. For COs, EVs were collected from conditioned media of 20-30 different COs in culture.

For the nanoparticle tracking analysis (NTA), fresh, unfrozen extracellular vesicle suspensions were diluted in PBS and analysed using a Particle Metrix ZetaView® PMX110-Z Nanoparticle Tracking Analyzer (Particle Metrix GmbH, Inning am Ammersee, Germany) equipped with a 520 nm laser. For each measurement, samples were introduced manually, the temperature was set to 24 °C, and two cycles were performed by scanning at 11 discrete positions in the cell channel and capturing 60 frames per position (video setting: high). The following recommended parameters were used for the measurement:

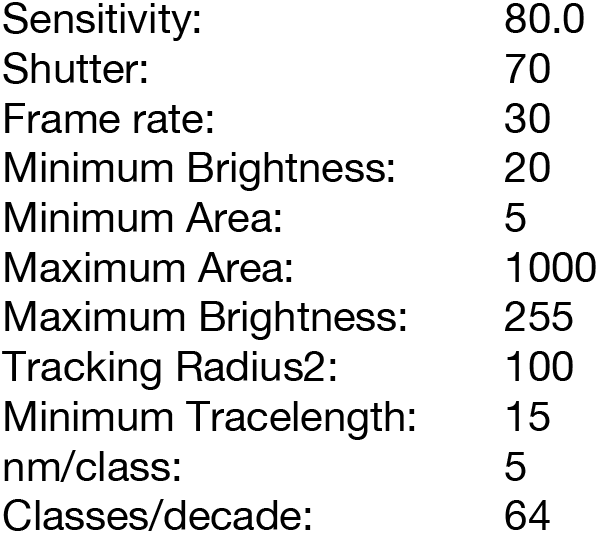

|  |  |
| --- | --- |
| Sensitivity: | 80.0 |
| Shutter: | 70 |
| Frame rate: | 30 |
| Minimum Brightness: | 20 |
| Minimum Area: | 5 |
| Maximum Area: | 1000 |
| Maximum Brightness: | 255 |
| Tracking Radius2: | 100 |
| Minimum Tracelength: | 15 |
| nm/class: | 5 |
| Classes/decade: | 64 |

After capture, the videos were analysed for particle size and concentration using the ZetaView Software 8.05.12 SP1.

For immune-electron microscopy, aliquots of extracellular vesicle suspensions were anayzed by Dr Ilkka Miinalainen at Biocenter Oulu / EM laboratory, Finland (Deun et al., 2020). Vesicles were deposited on Formvar carbon coated, glow-discharged grids and incubated in a blocking serum containing 1% BSA in PBS. CD81, CSTB primary antibodies and secondary gold conjugates (Zymed, San Francisco, CA, USA) were diluted in 1% BSA in PBS. The blocking efficiency was controlled by performing the labelling procedure in the absence of primary antibody.

### Dissociation of ventral and dorsal organoids for 2D cultures

2 months old ventral and dorsal d- and v-COs generated from control and EPM1 iPSCs, were dissociated as previously described (Di Matteo et al., 2020), with some modifications. Briefly COs were dissociated to single cells using Accutase (Sigma-Aldrich, A6964). Single cells were then plated onto Poly-L-ornithine (10 μg/ml) (Sigma-Aldrich, P4957)/Laminin (10 μg/ml) (Sigma-Aldrich, L2020)- coated coverslips with some modifications in wells of 24-well plates (Corning). Cells coming from a pool of two to three organoids were plated in 12 wells of 24-well plates with neural progenitor cells medium (NPC medium) (Gunhanlar et al., 2018).

### Immunohistochemistry

Cells dissociated from 2 months old COs were fixed using 4% PFA for 10 min and permeabilized with 0.3% Triton for 5 min. After fixation and permeabilization, cells were blocked with 0.1% Tween, 10% Normal Goat Serum (Biozol, VEC-S-1000). Primary and secondary antibodies were diluted in blocking solution (Table 1). Stained cells were analyzed using a Leica laser-scanning microscope.

**Table 1.** Immunostaining antibodies

| Primary antibodies |  |  |  |  |
| --- | --- | --- | --- | --- |
| Antibody | Host | Company | Catalogue No. | Dilution |
| TBR1 | Rabbit | Abcam | Ab31940 | 1:500 |
| CTIP2 | Rat | Abcam | Ab18465 | 1:500 |
| SATB2 | Mouse | Abcam | Ab51502 | 1:500 |
| GABA | Rabbit | Sigma | A2052 | 1:500 |
| CALB1 | Rabbit | Swang | CB-38a | 1:1000 |
| CALB2 | Rabbit | Swang | CRGP7 | 1:1000 |
| GAD67 | Mouse | Millipore | MAB5406 | 1:1000 |
| NKX2.1 | Mouse | Merk | MAB5460 | 1:500 |
| MEIS1/2/3 | Mouse | Santa Cruz Biotechnology | SC 101850 | 1:300 |
| KI67 | Rabbit | Abcam | AB15580 | 1:500 |
| GFP | Chicken | Rockland | 600-901-215 | 1:500 |
| Secondary antibodies (confocal microscopy) |  |  |  |  |
| Alexa Fluor® 488 Goat Anti-Rabbit IgG (H+L) | Goat | ThermoFisher Scientific | A11008 | 1:500 |
| Alexa Fluor® 488 Goat Anti-Chicken IgG (H+L) | Goat | ThermoFisher Scientific | A11039 | 1:500 |
| Alexa Fluor® 546 Goat Anti-Mouse IgG (H+L) | Goat | ThermoFisher Scientific | A21143 | 1:500 |
| Alexa Fluor® 647 Goat Anti-Mouse IgG (H+L) | Goat | ThermoFisher Scientific | A21242 | 1:500 |
| AlexaFluor647 Alexa Fluor® 647 Goat Anti-Rabbit IgG (H+L) | Goat | ThermoFisher Scientific | A21244 | 1:500 |
| Alexa Fluor® 546 Goat Anti-Rat IgG (H+L) | Goat | ThermoFisher Scientific | A11081 | 1:500 |

NPCs and neurons were fixed with 4% paraformaldehyde for 20 mins at room temperature, followed by three times 5 min washing with 1xPBS. Next, cells were blocked against unspecific binding and permeabilized in blocking buffer (10% Normal Goat Serum, 0,02% Triton-X in 1xPBS) for 1 hour. Primary antibodies diluted in blocking buffer were then added to the cells in the dilutions specified below and incubated overnight. On the second day, cells were washed five times for 5 mins each in PBS with 0,1% Tween (PBS-T), and then incubated for 2 hours in secondary antibodies raised against the host animal of the primary antibody. Secondary antibodies were diluted in blocking buffer and the dilutions used are listed below (Table 1). DAPI (4′,6-diamidino-2-phenylindole) was added as a nuclear counterstaining. Finally, cells were washed three times with PBS-T and mounted on object slides with Fluoromount-G (ThermoFisher Scientific, 00-4958-02).

### Imaging

Confocal stack images were obtained using a Leica SP8 confocal microscope based on a DMi8 stand (Leica Microsystems, Wetzlar, Germany), equipped with 20x/0.75 (oil), 40x/1.10 (water), and 63x/1.30 (glyc) objectives. Images were then processed using ImageJ (Schneider et al., 2012).

For the live imaging of neuron exocytosis, young neurons (1-2 weeks) from two control NPC lines and two patient NPC lines were transfected with the plasmid pCMV-Sport6-CD63-pHluorin (A gift from DM Pegtel (Addgene plasmid # 130901; http://n2t.net/addgene:130901; RRID:Addgene_130901) using the Lipofectamine™ 3000 Transfection Reagent (ThermoFisher Scientific, USA) as instructed in the protocol. 48h following the transfection, cells were imaged using a Leica TIRF system and a 100x/1.47 NA objective as previously described (Verweij et al., 2018b). The videos obtained were analyzed using the AMvBE (Analyzer of Multivesicular Body Exocytosis) macro previously developed (Verweij et al., 2018b) for ImageJ (Schneider et al., 2012).

### Silicon probe recordings in COs

For recordings of spontaneous spike activity in COs, we followed Di Matteo et al. (Di Matteo et al., 2024). A a 8-month-old CO was glued with a tiny drop (0.25 µl) of Histoacryl (B.Braun) on a polypropylene mesh that was mounted on a circular plastic frame. Afterwards, the mCO was incubated at 37oC for at least 30 minutes in carbogen gas (95% O2/5% CO2)-saturated ACSF consisting of (in mM): 121 NaCl, 4.2 KCl, 29 NaHCO3, 0.45 NaH2PO4, 20 Glucose, 0.5 Na2HPO4, 1.1 CaCl2 and 1 MgSO4. Subsequently, the CO was transferred with the holding device to the recording chamber and superfused with warm (37oC) carbogenated ACSF at a flow rate of 2.5 ml/min. For additional stabilization, a custom-made anchor was gently placed on the top of the hCO. Recordings were performed using 16-channel probes (Cambridge Neurotech, ASSY-1 E1), which were connected to a ME2100 system (Multichannel Systems). The headstage was mounted on a PatchStar micromanipulator (Scientifica). Recording data were high-pass filtered at 100 Hz, low-pass filtered at 4 kHz, digitized at 20 kHz and transferred via a MCS IFB interface board (Multichannel Systems) to a personal computer. Data were stored using the software Multichannel Experimenter (Multichannel Systems). Visually guided insertion of the silicon probe into COs was conducted using a SliceScope microscope (Scientifica) equipped with a 2.5x objective. Once neuronal activity has been detected, the probe was allowed to stabilize in the tissue for 5 minutes and afterwards activity was recorded for 5 minutes. For each condition, independent recording areas were used as biological replicates. Recordings were performed in at least three independent areas per CO, keeping a distance of at least 400 µm between two adjacent areas. Independent COs, each from at least two different batches were analyzed. Sample size: Ctrl#1 n=27; Ctrl#2 n=28; EPM1.1 n=28; EPM1.2 n=25.

### MEA recordings

After 30 days in culture, dorsal and ventral organoids from every genotype were dissociated to single cells as previously described (Di Matteo et al., 2020). The dorsal and ventral cell suspensions with the same genotype were then mixed at a 50:50 ratio and plated into the 24-well MEA plate (Multichannel System) previously coated with poly-L-ornithine/Laminin. Each well contained 12 gold electrodes (30 µm electrode diameter, 300 µm electrode spacing). The neuronal differentiation was sustained for the following 7 weeks by supplementing the complete Neurobasal medium with 20 ng/ml BDNF, 20 ng/ml GDNF, 1 μM cAMP, 200 μM ascorbic acid, and 2 μg/ml laminin.

The recording of spontaneous spike activity was conducted 8 weeks after dissociation, using the Multiwell-MEA-System (Multichannel Systems). 5 min recordings were performed at 37oC under a humidified gas atmosphere (95% O2/5% CO2). Data were high-pass filtered at 1 Hz, low-pass filtered at 3.5 kHz, digitized at 20 kHz and stored with the Multiwell Screen software (Multichannel Systems). For each condition, independent wells were analyzed as biological replicates. At least three independent wells for each of the three different batches were analyzed. Analysis was conducted with the Multiwell Analyzer software (Multichannel Systems). For each recording, the channel with the highest activity was used for the analysis.

Sample size in homo-genotype condition: Ctrl#1 n=12; EPM1.1 n=12; EPM1.2 n=12.

Sample size in hetero-genotype condition: Ctrl#1 n=5; EPM1.1 n=5; EPM1.2 n=5.

### Proteomic analysis

#### Sample preparation for mass spectrometry

Purified EVs, collected from the conditioned media of 20-30 COs in culture, and IVs, isolated from a pool of 5-7 COs, were lysed in RIPA buffer (150mM NaCl, 50mM Tris pH8, 0.1% DOC, 0.1% SDS, 0.1% NP40). 10 ug of protein for each sample was subjected to the modified FASP protocol (Wiśniewski et al., 2009). Briefly, the protein extract was loaded onto the centrifugal filter CO10 kDa (Merck Millipore, Darmstadt, Germany), and detergent were removed by washing five times with 8M Urea (Merck, Darmstadt, Germany) 50mM Tris (Sigma-Aldrich, USA) buffer. Proteins were reduced by adding 5mM dithiothreitol (DTT) (Bio-Rad, Canada) at 37degrees C for 1 hour in the dark. To remove the excess of DTT, the protein sample was washed three times with 8M Urea, 50mM Tris. Subsequently protein thiol groups were blocked with 10mM iodoacetamide (Sigma-Aldrich, USA) at RT for 45 min. Before proceeding with the enzymatic digestion, urea was removed by washing the protein suspension three times with 50mM ammonium bicarbonate (Sigma-Aldrich, Spain). Proteins were digested first by Lys-C (Promega, USA) at RT for 2 hours, then by trypsin (Premium Grade, MS Approved, SERVA, Heidelberg, Germany) at RT, overnight, both enzymes were added at an enzyme-protein ratio of 1:50 (w/w). Peptides were recovered by centrifugation followed by two additional washes with 50mM ammonium bicarbonate and 0.5M NaCl (Sigma-Aldrich, Swisserland). The two filtrates were combined, the recovered peptides were lyophilized under vacuum. Dried tryptic peptides were desalted using C18-tips (Thermo Scientific, Pierce, USA), following the manufacture instructions. Briefly, the peptides dissolved in 0.1%(v/v) formic acid (Thermo scientific, USA) were loaded onto the C18-tip and washed 10 times with 0.1 % (v/v) formic acid, subsequently the peptides were eluted by 95% (v/v) acetonitrile (Merck, Darmstadt, Germany), 0.1% (v/v) formic acid. The desalted peptides were lyophilized under vacuum. The purified peptides were reconstituted in 0.1% (v/v) formic acid for LC-MS/MS analysis.

#### MS data acquisition

Desalted peptides were loaded onto a 25 cm, 75 µm ID C18 column with integrated nanospray emitter (Odyssey/Aurora, ionopticks, Melbourne) via the autosampler of the Thermo Easy-nLC 1000 (Thermo Fisher Scientific) at 60 °C. Eluting peptides were directly sprayed onto the timsTOF Pro (Bruker Daltonics). Peptides were loaded in buffer A (0.1% (v/v) formic acid) at 400 nl/min and percentage of buffer B (80% acetonitril, 0.1% formic acid) was ramped from 5% to 25% over 90 minutes followed by a ramp to 35% over 30 minutes then 58% over the next 5 minutes, 95% over the next 5 minutes and maintained at 95% for another 5 minutes. Data acquisition on the timsTOF Pro was performed using timsControl. The mass spectrometer was operated in data-dependent PASEF mode with one survey TIMS-MS and ten PASEF MS/MS scans per acquisition cycle. Analysis was performed in a mass scan range from 100-1700 m/z and an ion mobility range from 1/K0 = 0.85 Vs cm-2 to 1.30 Vs cm-2 using equal ion accumulation and ramp time in the dual TIMS analyzer of 100 ms each at a spectra rate of 9.43 Hz. Suitable precursor ions for MS/MS analysis were isolated in a window of 2 Th for m/z < 700 and 3 Th for m/z > 700 by rapidly switching the quadrupole position in sync with the elution of precursors from the TIMS device. The collision energy was lowered as a function of ion mobility, starting from 45 eV for 1/K0 = 1.3 Vs cm-2 to 27eV for 0.85 Vs cm-2. Collision energies were interpolated linear between these two 1/K0 values and kept constant above or below these base points. Singly charged precursor ions were excluded with a polygon filter mask and further m/z and ion mobility information was used for ‘dynamic exclusion’ to avoid re-sequencing of precursors that reached a ‘target value’ of 14500 a.u. The ion mobility dimension was calibrated linearly using three ions from the Agilent ESI LC/MS tuning mix (m/z, 1/K0: 622.0289, 0.9848 Vs cm-2; 922.0097 Vs cm-2, 1.1895 Vs cm-2; 1221.9906 Vs cm-2, 1.3820 Vs cm-2).

#### Raw data analysis of MS measurements

Raw data were processed using the MaxQuant computational platform (version 1.6.17.0) (Tyanova et al., 2016) with standard settings applied for ion mobility data (Prianichnikov et al., 2020). Shortly, the peak list was searched against the Uniprot database of Human database (75069 entries, downloaded in July 2020) with an allowed precursor mass deviation of 10 ppm and an allowed fragment mass deviation of 20 ppm. MaxQuant by default enables individual peptide mass tolerances, which was used in the search. Cysteine carbamidomethylation was set as static modification, and methionine oxidation, deamidation and N-terminal acetylation as variable modifications. The match-between-run option was enabled, and proteins were quantified across samples using the label-free quantification algorithm in MaxQuant generating label-free quantification (LFQ) intensities.

#### Bioinformatic analysis

For the proteomic characterization in EVs, 5489 proteins were quantified. Proteins that were detected with more than 1 peptide and consistently detected in 2 of the 3 technical replicates per each condition were retained. Downstream analysis was performed using Perseus (https://maxquant.net/perseus/). The LFQs values were log2-transformed. Missing values were replaced from normal distribution using the Perseus Imputation function (shift of 1.8 units of the standard deviation and a width of 0.3). Differentially expression (DEP) analysis was performed on the imputed data using t-Test in the Volcano plot function of Perseus. Proteins with FDR-corrected q-value < 0.05 and log2 fold change values (log2FC) ≥ 1 and ≤ -1 were considered as differentially expressed.

### Single-cell RNA-sequencing sample preparation

Single-cell dissociation was performed on five 40 days old-patterned spheroids randomly selected for each pattern condition. Single cells were dissociated using StemPro Accutase Cell Dissociation Reagent (Life Technologies), filtered through 30 uM and 20 uM filters (Miltenyi Biotec) and cleaned of debris using a Percoll (Sigma, P1644) gradient. Single cells were resuspended in ice-cold Phosphate-Buffered Saline (PBS) supplemented with 0.04% Bovine Serum Albumin at a concentration of 1000 cells per ul. Single cells were loaded onto a Chromium Next GEM Single Cell 3^’
s^ chip (Chromium Next GEM Chip G Single Cell Kit, 16 rxns 10XGenomics PN-1000127) with the Chromium Next GEM Single Cell 3′ GEM, Library & Gel Bead Kit v3.1 (Chromium Next GEM Single Cell 3′ GEM, Library & Gel Bead Kit v3.1, 4 rxns 10xGenomics PN-1000128) and cDNA libraries were generated with the Single Index Kit T Set A, 96 rxns (10xGenomics PN-1000213) according to the manufacturer’s instructions. Libraries were sequenced using Illumina NovaSeq6000 in 28/8/91bp mode (SP flowcell), quality control and UMI counting were performed by the Max-Planck für molekulare Genetik (Germany).

#### Single-cell RNA sequencing data analysis

Raw sequencing data were processed using Cell Ranger v6.0.0 (10x Genomics), a software suite for analyzing single-cell RNA sequencing (scRNA-seq) data. The count function within Cell Ranger was utilized for read alignment, quantification of gene expression, and generation of a gene-cell matrix. The analysis was conducted on a high-performance computing cluster, using 250 GB of memory and 32 CPU cores on a single node. The human reference genome, GRCh38 version 1.2.0, was used for reads alignment and transcriptome generation, with intronic reads included to capture nascent transcripts, and secondary alignments excluded.

Quality control (QC) and preprocessing of single-cell RNA-seq data were performed using Scanpy (Wolf et al., 2018). Cells with low quality, such as those with a high proportion of mitochondrial reads or an abnormally low number of genes expressed, were removed to ensure the retention of high-quality data for further analysis.

To remove low-quality cells from the dataset, an outlier detection strategy was implemented based on key quality metrics. Cells were filtered using thresholds based on the median and median absolute deviation (MAD) of these metrics. Specifically, outliers were defined as cells where any of the following conditions were met: the total counts per cell (log1p_total_counts) deviated more than 5 MADs from the median, the number of genes expressed per cell (log1p_n_genes_by_counts) deviated more than 5 MADs from the median, or the percentage of counts in the top 20 most expressed genes (pct_counts_in_top_20_genes) deviated more than 5 MADs from the median.

Additionally, mitochondrial content was used to further identify and remove low-quality cells. Cells were considered outliers if the percentage of mitochondrial gene expression (pct_counts_mt) deviated more than 3 MADs from the median or exceeded a fixed threshold of 8%.

To identify and remove potential doublets from the dataset, the scDblFinder algorithm was used within the R environment, integrated into the Python workflow using rpy2. The raw count matrix was transferred from the Scanpy object into the R environment, where the scDblFinder package was used to classify cells as doublets or singlets. Specifically, the SingleCellExperiment framework was used to store the expression data, and doublets were identified using scDblFinder with a random seed set for reproducibility. Following this process, each cell was assigned a doublet score and a classification as either a doublet or singlet. Cells classified as doublets were excluded from the dataset before proceeding with downstream analyses, ensuring that only singlet cells were retained for further clustering and gene expression analysis.

Normalization was performed using the scran method to account for variations in library size. Size factors were computed for each cell using the computeSumFactors function, leveraging preliminary clustering to improve accuracy. The count matrix was then divided by these size factors, and a log1p transformation was applied to the normalized data. This scran-normalized data was stored in the adata.layers for downstream analyses. Highly variable genes were selected for downstream analysis using Scanpy’s highly_variable_genes function. Principal Component Analysis (PCA) was applied to reduce the dimensionality of the dataset, and the top 30 principal components were used for neighborhood graph construction. To cluster the cells, the Leiden algorithm (Traag et al., 2019) was employed using a resolution of 1. The neighborhood graph was constructed using the pp.neighbors function, and clustering was performed using tl.leiden. To visualize the clusters, Uniform Manifold Approximation and Projection (UMAP) was performed with the tl.umap function in Scanpy. Clusters were visualized in two-dimensional space, and marker genes were used to assign cell types to the clusters. Differential gene expression analysis was performed between clusters using the Wilcoxon rank-sum test to identify potential marker genes for each cell population. Cell types were also annotated using CellTypist. The pretrained model developing_Human_Brain.pkl was applied for automated cell type prediction, as described by Xu et al. (Xu et al., 2023)and Domínguez Conde et al. (Domínguez Conde et al., 2022). This automated annotation was run in parallel with marker gene-based manual annotation to ensure accuracy and consistency in cell type identification. Cells identified as fibroblasts through the automated annotation with CellTypist were excluded from the dataset to focus on other relevant cell populations. We retained a total of 10,972 cells for the ventral organoid experiments and 10,965 cells for the mosaic experiment. These high-quality cells were used for all subsequent analyses. Following this, batch effects were corrected, and datasets were integrated using Scanorama to ensure a harmonized view across different samples. The integration was performed to account for technical variations while preserving the biological variance of interest.

Pseudobulk RNA-seq analysis was performed by aggregating the counts of cells belonging to specific cell types into pseudo-replicates. Cells annotated as progenitors were selected, and counts from these cells were summed across pseudo-replicates. The resulting pseudobulk data were analyzed using the DESeq2 pipeline via the pyDESeq2 package. Differential expression analysis was conducted between different conditions, and the results were visualized using volcano plots to identify significantly up- and down-regulated genes. Specific gene sets of interest, such as those involved in the Wnt signaling pathway and dorsoventral patterning, were highlighted in the analysis.

All analyses were performed using Python version 3.9.18. The following packages were used for the analysis: Scanpy version 1.9.6, Scanorama version 1.7.4, NumPy version 1.26.2, Pandas version 2.1.1, Matplotlib version 3.7.1

### Bulk-RNA-sequencing

RNA-seq was performed on 10ng of total RNA collected from 3 independent wells of NPCs from a 24well plate. NPCs were not treated with EVs or treated for 12h with EVs collected by ultracentrifugation from 25 ml of conditioned medium collected from 28 to 37 days in culture COs (control ventral, EPM1 ventral). NPCs were lysed in 1ml Trizol (Qiagen)/well and RNA was isolated employing RNA Clean & Concentrator kit (Zymo Research) including digestion of remaining genomic DNA according to producer’s guidelines. RNA was further processed according to (Cernilogar et al., 2019). Briefly, cDNA synthesis was performed with SMART-Seq v4 Ultra Low Input RNA Kit (Clontech cat. 634888) according to the manufacturer’s instruction. cDNA was fragmented to an average size of 200–500 bp in a Covaris S220 device (5 min; 4°C; PP 175; DF 10; CB 200). Fragmented cDNA was used as input for library preparation with MicroPlex Library Preparation Kit v2 (Diagenode, cat. C05010012) and processed according to the manufacturer’s instruction. Libraries were quality controlled by Qubit and Agilent DNA Bioanalyzer analysis. Deep sequencing was performed on a HiSeq 1500 system according to the standard Illumina protocol for 50 bp paired-end reads with v3 sequencing reagents.

#### Bulk RNA seq analysis

Paired end reads were aligned to the human genome version GRCh38 using STAR v2.6.1d (Dobin et al., 2013) with default options “--runThreadN 32 -- quantMode TranscriptomeSAM GeneCounts -- outSAMtype BAM SortedByCoordinate”. Reads-per-gene counts were imported in R v4.1.0.

Expression data from BulkRNA seq samples were loaded on Rstudio, focusing on transcript-per-million (TPM) values. Data normalization was performed by calculating Z-scores across expression values, facilitating comparability between samples, and rows with missing values were excluded to ensure data integrity. Hierarchical clustering was employed to uncover patterns in expression profiles. Clusters were visualized on a heatmap, with both rows (genes) and columns (samples) grouped based on similarity. Gene clusters were extracted, and a final list of genes within each cluster was compiled and exported for further analyses. The R packages ComplexHeatmap (Gu et al., 2016), circlize (Gu et al., 2014), colorspace (Zeileis et al., 2009), GetoptLong (https://cran.r-project.org/web/packages/GetoptLong/), and ggpattern (https://github.com/coolbutuseless/ggpattern) were used for heatmap visualization, clustering, and customization. dendsort (Sakai et al., 2014) refined hierarchical clustering, while RColorBrewer (https://cran.r-project.org/web/packages/RColorBrewer/) and colorspace (Zeileis et al., 2009)managed color gradients. pheatmap (https://cran.r-project.org/web/packages/pheatmap/) enabled targeted gene display, and write.table exported clustered gene lists.

#### DATA AND CODE AVAILABILITY

Bulk RNA Seq data and scRNAseq data have been deposited in the GEO database.

#### -Enrichment analysis

GO term analysis of differentially expressed proteins was tested using the FUMA algorithm (Watanabe et al., 2017) by inserting the DE protein lists into the GENE2FUNC software (FDR<0.05) (https://fuma.ctglab.nl/) or with STRING (https://string-db.org).

## Acknowledgments

we thank I.Miinalainen, Maik Ködel, G. Masserdotti, M. Ianni, Natalia Abate for technical help and critical discussion. We thank the International Human Frontier Science Program Organization (grant number 5627cc47) and the Alexander von Humboldt Foundation for supporting MVP.

## Author contributions

Conceptualization: SC, RDG.Methodology: RDG, AF, MVP, FP, AS, FDM, ZB, EF, GM, MGP, FMC. Investigation: SC, RDG, MVP, AF, FP, FDM, EF, GM, MGP, FMC. Visualization: SC, RDG, AF, MVP, FP. Funding acquisition: SC, RDG.Supervision: SC, RDG.Writing – original draft: RDG, MVP; review & editing: SC, RDG, MVP, AF.

## Supplementary figures

**Supplementary Fig. 1.**
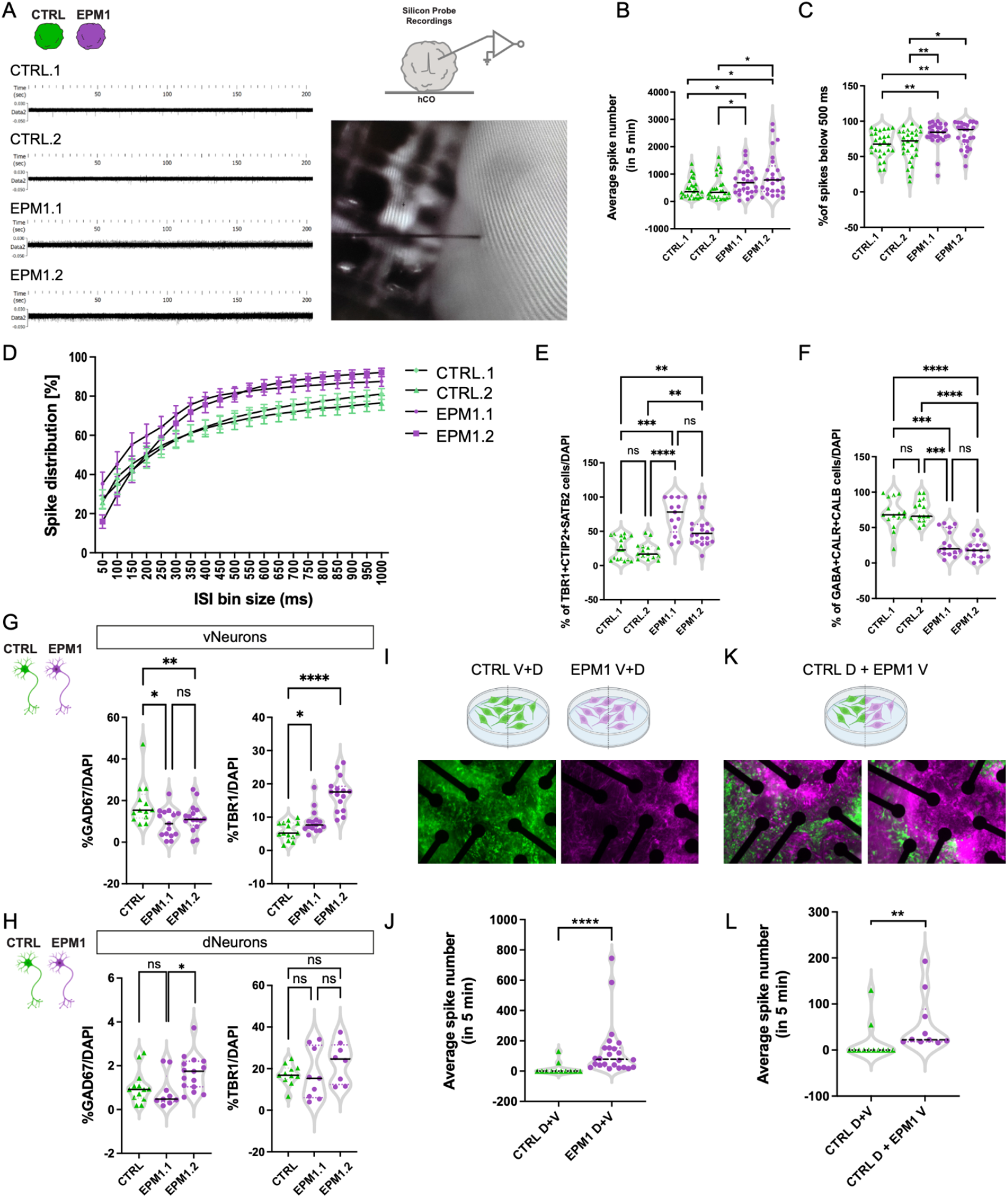
Comprehensive analysis of electrophysiological activity and neuronal composition in EPM1 cerebral organoids. (A) Left, Representative electrophysiological traces of COs from two control iPSC lines (CTRL.1 and CTRL.2) and two patient iPSC lines (EPM1.1 and EPM1.2). Right, scheme and representative image of the silicone probe used for electrophysiological measurements in COs. (B) Average spike number detected within 5 min for COs from two control iPSC lines (CTRL.1 and CTRL.2) and two patient iPSC lines (EPM1.1 and EPM1.2). n = 25-28 per condition. Statistical significance was determined using the a Mann-Whitney test, *p<0.05. (C) Percentage of spikes above 500 ms detected in COs from two control iPSC lines (CTRL.1 and CTRL.2) and two patient iPSC lines (EPM1.1 and EPM1.2). n = 25-28 per condition. Statistical significance was determined using the a Mann-Whitney test, *p<0.05, **p<0.01. (D) Percentage of spike distribution at interspike interval bins ranging from 50 to 1000 ms for COs from two control iPSC lines (CTRL.1 and CTRL.2) and two patient iPSC lines (EPM1.1 and EPM1.2). (E) Quantification of the percentage of cells positive for TBR1/CTIP2/SATB2 in neural cultures from two control iPSC lines (CTRL.1 and CTRL.2) and two patient iPSC lines (EPM1.1 and EPM1.2). n = 12-18 per condition. Statistical significance was determined using the a Kruskal-Wallis test, **p<0.01, ***p<0.001, ****p<0.0001. (F) Quantification of the percentage of cells positive for GABA/ CALB1/ CALB2 in neural cultures from two control iPSC lines (CTRL.1 and CTRL.2) and two patient iPSC lines (EPM1.1 and EPM1.2). n = 15 per condition. Statistical significance was determined using the a Kruskal-Wallis test, ***p<0.001, ****p<0.0001. (G) Quantification of GAD67-positive cells/DAPI and TBR1-positive cells/DAPI in vNeurons from one control iPSC lines (CTRL.1) and two patient iPSC lines (EPM1.1 and EPM1.2). n = 13-16 per condition. Statistical significance was determined using the a Kruskal-Wallis test, *p<0.05. (H) Quantification of GAD67-positive cells/DAPI and TBR1-positive cells/DAPI in dNeurons from one control iPSC lines (CTRL.1) and two patient iPSC lines (EPM1.1 and EPM1.2). n = 8-15 per condition. Statistical significance was determined using the a Kruskal-Wallis test, **p<0.01, ****p<0.0001. (I) Schematic of single cultures of ventral (V) and dorsal (D) progenitors dissociated from CTRL and EPM1 COs (top), and representative images of CTRL (green) and EPM1 (magenta) cells seeded separately in MEA chip. (J) Average spike number detected within 5 min for single cultures of ventral (V) and dorsal (D) progenitors dissociated from CTRL and EPM1 COs using MEA. (K) Schematic of mixed neuronal culture from ventral (V) EPM1 and dorsal (D) CTRL progenitors dissociated from COs (top), and representative images of CTRL (green) and EPM1 (magenta) cells seeded together in MEA chip. (L) Average spike number detected within 5 min for mixed neuronal culture from ventral (V) EPM1 and dorsal (D) CTRL progenitors dissociated from COs using MEA. Abbreviations: milliseconds (ms); minutes (min); Multi-electrode Array (MEA)

**Supplementary Fig. 2.**
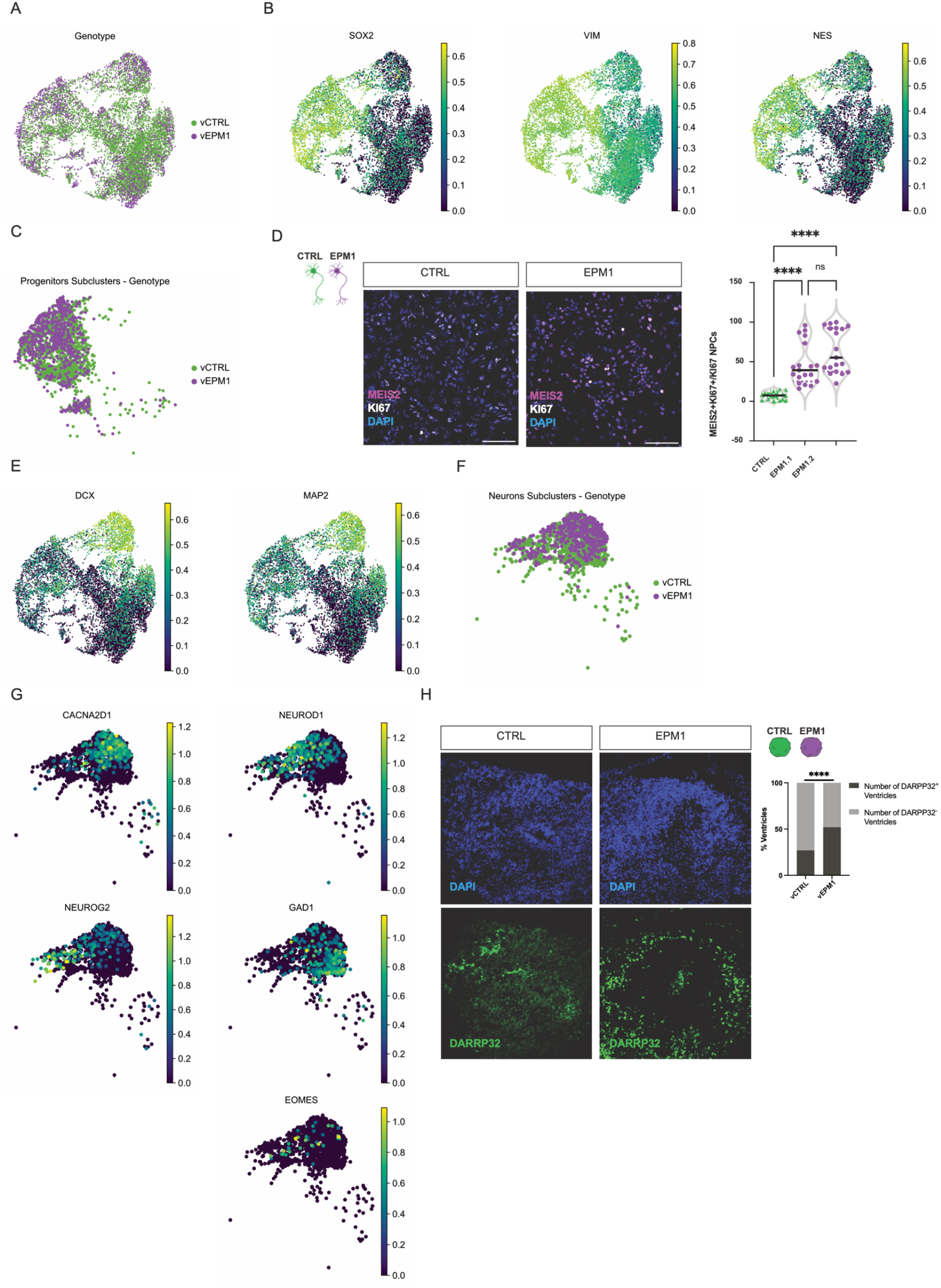
Supplementary scRNA-seq analysis of progenitor and mature neuron markers, with additional validations in ventral control (vCTRL-COs) and EPM1 cerebral organoids (vEPM1-COs). (A) UMAP visualization of scRNA-seq data from CTRL and EPM1 vCOs, clustered by cell genotype. (B) UMAP visualization of SOX2, VIM and NES expression. (C) UMAP visualization showing the distribution of vCTRL and vEPM1 cell within the Progenitors Subclusters. (D) Representative images of MEIS2 (Magenta) and KI67 (White) immunostaining in CTRL and EPM1 neural cultures (left), and quantification of MEIS2+/KI67+ NPCs in CTRL and two patient-derived (EPM1.1 and EPM1.2) neural cultures (right). (E) UMAP visualization of DCX and MAP2 expression across cells. (F) UMAP visualization showing the distribution of vCTRL and vEPM1 cells within the Mature Neuron Subclusters. (G) UMAP visualization highlighting the expression of excitatory neuronal markers (CACNA2D1, NEUROD1), inhibitory neuronal markers (GAD1), and dorsal telencephalon markers (NEUROG2, EOMES). (H) Representative images of DARRP32 immunostaining in CTRL and EPM1 vCOs, and quantification of DARRP32+ ventricles in CTRL and EPM1 COs.

**Supplementary Fig. 3.**
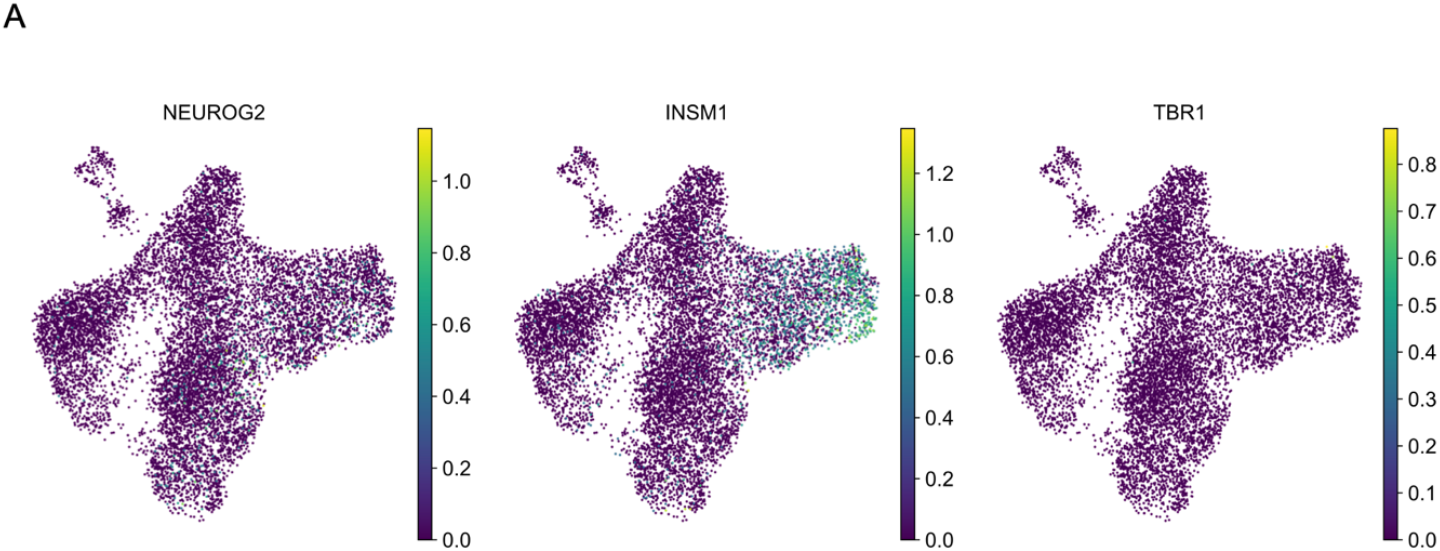
Supplementary scRNA-seq analysis of mature neuron markers in ventral hybrid control (hCTRL-COs) and EPM1 cerebral organoids (hEPM1-COs). (A) UMAP visualization of the expression of excitatory neuronal markers (NEUROD1, CACNA2D1), inhibitory neuronal (DLX2, GAD1) and dorsal telencephalon markers (NEUROG2, TBR1).

**Supplementary Fig. 4.**
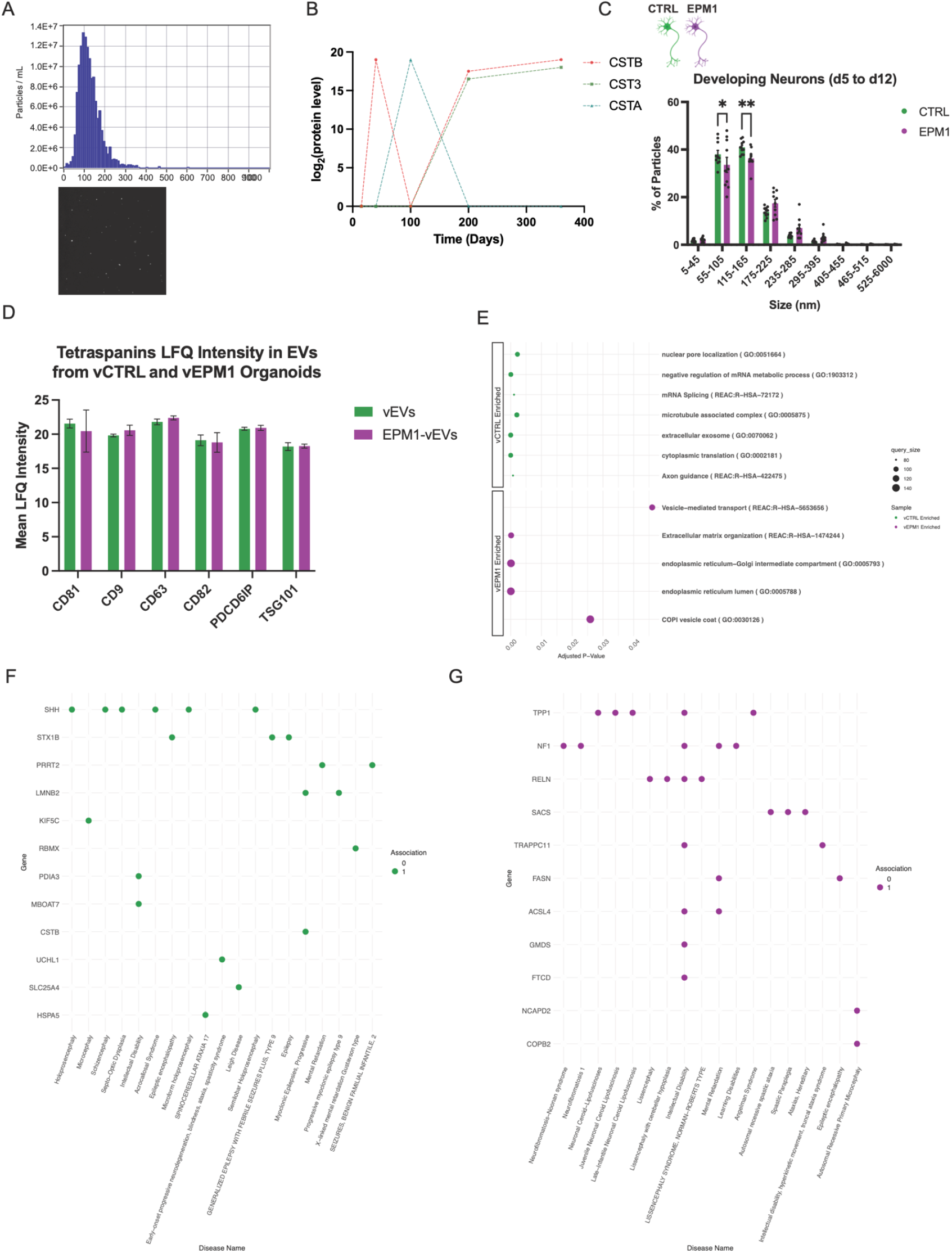
Nanoparticle Tracking Analysis (NTA) and disease association of EPM1 EVs. (A) Example distribution plot and image illustrating EV quantification via NTA. (B) Expression levels of CSTB, CSTA and CST3 protein levels in extracellular vesicles derived from COs at different developmental stages (expressed in days). (C) Percentage of particles detected in NTA of EVs isolated from CTRL and EPM1 NPCs and differentiating neurons (5 to 12 days of differentiation). (D) Expression levels of Tetraspanins protein levels in extracellular vesicles derived from CTRL and EPM1 vCOs. (E) Gene Ontology (GO) terms enriched in EV proteins from vCTRL (top) and vEPM1 (bottom) samples. (F) List of proteins enriched in EPM1-vEVs that are associated with brain diseases, as identified using DisGeNET. (G) List of proteins enriched in CTRL-vEVs that are associated with brain diseases, as identified using DisGeNET.

